# Neuregulin-1 exerts molecular control over axolotl lung regeneration through ErbB family receptors

**DOI:** 10.1101/258517

**Authors:** Tyler B Jensen, Peter Giunta, Natalie Grace Schulz, Yaa Kyeremateng, Hilary Wong, Adeleso Adesina, James R Monaghan

## Abstract

The induction of new lung tissue after disease or trauma has the potential to save lives and transform patient outcomes. Ambystoma mexicanum, the axolotl salamander, is a classic model organism used to study vertebrate regeneration, primarily after limb amputation. While it is hypothesized that axolotls regenerate all of their tissues, exploration of lung regeneration has not been performed until now. Proliferation after lung injury was observed to be a global response, suggesting that regeneration utilizes a compensatory mechanism, in contrast to limb regeneration’s epimorphic response. ErbB signaling is crucial for the proliferative response during lung regeneration, likely through the ErbB2:ErbB4 receptor heterodimer. ErbB4 mRNA was found to be highly upregulated at both one and three weeks post amputation. Neuregulin-1p (NRG1) can induce proliferation in the lung and likely exerts molecular control over lung regeneration. Inhibition of ErbB2 was sufficient to both block regeneration and the proliferative response observed after NRG1 treatment.

## BACKGROUND

Each year there are 200,000 cases of acute respiratory distress syndrome (ARDS), a chronic condition that is the result of an acute lung injury (ALI)^1^. There are few therapeutic interventions that may be performed for patients, and physicians must rely on mechanical ventilation and assistive oxygen therapy until symptoms diminish and the patient can recover normal lung function^2^. Regenerative therapy after ALI could provide an alternative treatment plan for patients who cannot respire effectively. Molecular approaches to speeding up the healing process in the lung, as well as regenerating lost tissue, are vital to these patients’ health and survival. While there has been evidence of compensatory growth in murine, canine, and human lungs, restoration of lung surface area and tissue mass can take extensive time^3,4^. In humans, there is evidence that a 77% volume increase is possible after pneumonectomy, over the course of 15 years^5^. There are stem cells residing in the lung that may replenish the tissue, but their proliferation to regenerate the lungs is slow^6^. Mechanisms through which these stem cells may be activated and induced to replenish lost pulmonary tissue have yet to be determined, but hold therapeutic potential for patients after acute lung injury and loss of pulmonary volume and mass^7^.

The role of epidermal growth factor receptor family (ErbB) ligands and receptors in lung regeneration has been a subject of considerable research as of late ^8^. There are four known ErbB family members, named 1-4, with ErbB1 also referred to as EGFR, and each has distinct ligand binding regions and intracellular pathways that they can activate ^9^. Among the family members, ErbB2 has been shown to have no ability to bind ligands, gaining specificity through its heterodimer binding partner ^10^. Each of these receptors, as a receptor tyrosine kinase, are embedded independently in the cellular membrane until activated. Once activated by an extracellular ligand, receptors homo/heterodimerize and conduct their respective signaling cascade ^11^.

No pneumocyte growth factor has been established, though there are purportedly several candidates that have been found^12^. Neuregulin-1β (NRG1) has been hypothesized as a candidate molecule, and it has been shown *in vitro* in human lung epithelial cells that NRG1 can induce proliferation via the JAK-STAT pathway^13^. NRG1 activated ErbB4 serves as a dedicated receptor for the Hippo-Yap pathway, and the Hippo/Yap pathway has been recently indicated in the promotion and control of epithelial proliferation in the adult rat lung^14,15^. Activated ErbB4 is able to travel to the nucleus, enhancing the transcriptional activity of Yap and triggering proliferation^16^. Interleukin-1β has been shown to induce shedding of NRG1, potentially providing an indication of a mechanism through which NRG1 may give rise to proliferation after injury^17^.

The axolotl salamander is the oldest regenerative laboratory species, and has been used as a model of regeneration for hundreds of years, though the focus has been on limb and tail regeneration ^18,19^. Little research has been performed^20^ investigating the mechanisms of organ regeneration in this species, and there is great potential for new discoveries. The limb, after amputation, forms a blastema, which restores the limb exactly as it was before injury (epimorphic) ^21^. In contrast to this regeneration, most known regenerative responses of organs in nature take place through a compensatory mechanism, growing the remaining tissue larger, while not restoring the exact morphology of the organ ^22^. Understanding how these processes are controlled, as well as uncovering the master regulators of regeneration is vital to our fight against human disease. In this study, we perform the first investigation of lung regeneration in this species, and seek to understand the role of NRG1 and ErbB family signaling in the regenerative response observed. Using research performed on lung tissue in mammalian models, as well as what we know of limb regeneration, we propose a role for the ErbB family receptors. We observed that NRG1β, a ligand for ErbB4, was important during lung cell proliferation *in vivo*, and interrogated the role of this molecule and the ErbB2:ErbB4 heterodimer.

## RESULTS

### Tissue repair after injury

The axolotl lung comprises of alveolar folds, with arches of smooth muscle and ciliated cells cresting the folds ^23^. There is a single type of pneumocyte in the lung, contrasting mammalian species, which have two, with the type II pneumocytes serving as the stem cell niche of the lung^23,24^. Of the two mammalian pneumocytes, the axolotl’s would appear most similar to type II pneumocytes, and are likely the source of any proliferative replenishment. To identify pneumocytes in the axolotl lung and lay the foundation for our further study of this organ system, lung tissue was examined using histological staining for alkaline phosphatase activity and immunohistochemical staining for the type II keratin, Keratin 7. We found that both pneumocytes and epithelial tissues on the surface of the lung stained positive for Krt7. In contrast, alkaline phosphatase staining was strong primarily in pneumocytes lining the respiratory epithelium of the lung. The use of these complementary stains allowed us to histologically identify specific epithelial layers of the mature lung, aiding in our understanding of the lung structure during following experiments (Supplement material).

Regeneration after pneumonectomy of the distal third of the lung was examined to understand the axolotl’s healing response (Fig. 1A modified from Farkas and Monaghan 2016). Tissues were collected at three days post amputation (dpa), 7 dpa, and without injury to investigate the early healing response (n=4) (Fig. 1B). Masson’s trichrome staining showed that lung epithelium closed rapidly by one-week post amputation (wpa). At 3 dpa there was a significant clot on the end of the tissue, and there was inflammation disrupting the normal lung structure. By 1 wpa the inflammation had abated, and lungs were histologically similar to uninjured lungs. There was no additional collagen staining observed in the lung after injury suggesting that the wound does not form a scar after injury, contrary to mammalian lung lacerations ^25^.

**Figure 1:**
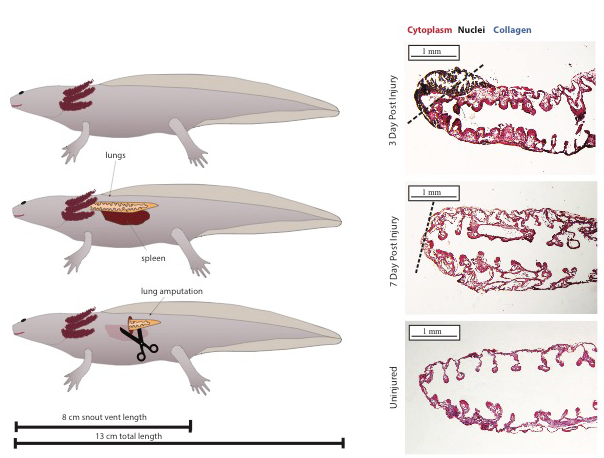
Surgery and wound closure after lung amputation. Fig. 1. (A) Representative image of model organism (axolotl) and the surgery performed, including localization of relevant organs and animal size at the time of experimentation and surgery. Left lung was surgically injured by amputation of approximately a third, and the right lung was left untouched. (B) Representative trichrome stains of the lung at 3 dpa, 7 dpa, and in control. It can be seen that the lung tissue has closed after one week, and rapidly regains normal appearance after injury. At three days the clot and inflammatory response can be seen in the injured lung. Dashed lines indicate plane of amputation. No additional collagen deposition was observed after lung injury, and would indicate a robust regenerative response.

### Proliferative Response and Tissue Recovery

We first sought to observe the localization of proliferation during lung regeneration (Fig. 2A). Regeneration in limbs and tails occurs by the formation of a blastema, and the proliferative response is observed only within close proximity to the wound site ^26^. We sought to determine whether the lung would form blastema tissue like that seen in appendages, or whether it would undergo compensatory regeneration as seen in other species ^27^. Lung regeneration was compared among uninjured (control), one, three, and six wpa (n=4 per time point) using bromodeoxyridine (BrdU) DNA synthesis analysis (Fig. 2B,C). Animals were pulsed intra-peritoneal (IP) with BrdU and collected. Proliferation was measured in the injured lung and the contralateral lung, comparing relative distal and proximal lung tissue proliferation, normalized to DAPI nuclear staining counts. There were no significant differences in proliferative responses between the locations measured, whether in tissue close to the airway or in the distal alveolar folds. Proliferation counts were observed to be equivalent in both injured and contralateral lung tissue. Lung proliferation was seen to be a systemic response, with BrdU-labelled cells increasing globally throughout the lung tissue. This would indicate that regeneration was a compensatory response, in which the entire lung grew larger to compensate for the missing tissue removed by amputation. Dividing cell types included mesenchymal cells, ciliated cells, and epithelial pneumocytes. Proliferation peaked at three weeks post-amputation, providing a time point to target for further proliferation assays.

**Figure 2:**
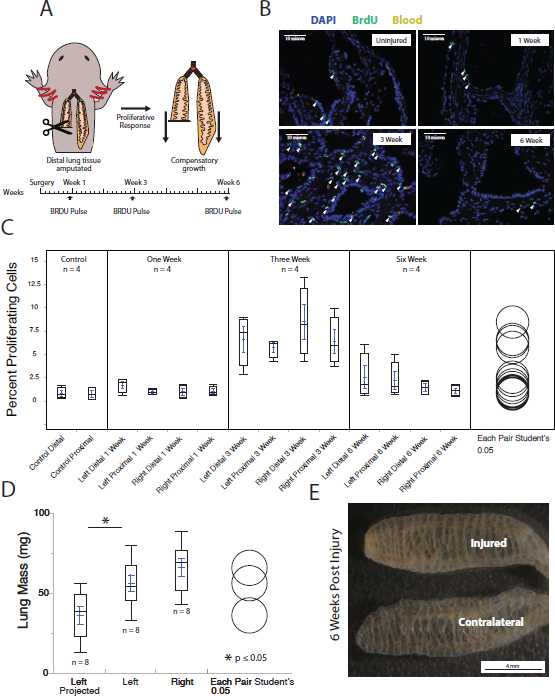
Compensatory Regenerative Response in the Lung after Amputation. Fig. 2. (A) Representative experimental outline and timeline of surgeries and collections with relative timing of BrdU pulsing. Left lung was surgically injured in this experiment while right lung was left intact. (B) Representative immunofluorescent images showing cell proliferation by anti-BrdU (green) antibody staining in control, 1 wpa, 3 wpa, and 6 wpa. BrdU(+) cells are indicated with white arrowheads. (C) Percent of BrdU(+) cells at control, 1 wpa, 3 wpa, and 6 wpa, showing a significant increase in proliferation at 3 weeks compared to control (p < 0.05) with no significant difference between proximal regions (close to airway) and distal regions (far from airway) or amongst injured and contralateral lungs. (D) Lung mass after regeneration of the left lung (injured) relative to the right lung (contralateral). Left projected was calculated by measuring the amount of tissue removed during surgery normalized to body weight, and was compared to actual lung mass at eight weeks after injury. Tissue was found to have regenerated faster than expected, thus showing increased growth rate in the left lung (Two tailed T-Test with unequal variance p = 0.02). (E) Representative image of left (injured) and right (contralateral) lungs at six wpa.

To measure the amount of tissue recovered after lung injury, the distal third of the lung was amputated, and the removed tissue was weighed. (Fig. 2D,E). While the lung length appeared impaired in the injured lung, volume appeared enlarged, and lung mass was recovered significantly. Over the course of eight weeks the injured lung caught up to the right lung as the pulmonary tissue responded to the injury. This suggests that the pulmonary tissue was able to recover significant volume in response to amputation, and recovers very quickly when compared to injuries in mammals.

### qPCR of one week and three week tissues after injury

ErbB family activity has been shown to be responsible for proliferative responses to injury in many tissue types, including hepatic^28^, cardiac^29^, and pulmonary^30^ tissues. Specifically, in pneumocytes, there are multiple indications that this family of receptors is critical to proliferation^30,31^. Additionally, YAP dysregulation has been shown to be mitogenic to lung epithelial tissue^32^. YAP activation is downstream of the ErbB receptor family, specifically through ErbB4, and could provide indications of an underlying mechanism through which these receptors regulate the intracellular effectors of proliferation^33^.

We examined ErbB1- 4 expression by qPCR analysis one (n=4) and three (n=4) wpa to discover the role of each receptor in controlling the proliferation observed in the pulmonary tissue (Fig. 3A,B). It was found that ErbB4 was highly upregulated, with ErbB1 also enriched. There was no significant enrichment of ErbB2 or ErbB3 at these time points. In mammals, ErbB2/ErbB4/NRG signaling is implicated in activation of proliferative genes, and can serve multiple roles in lung tissue^30^, while ErbB1 activation is implicated in promoting differentiation and enhancing pneumocyte maturation^34^. Downstream effects of activation of ErbB4 can include shuttling of STAT5 to the nucleus^35^, and activation and enhancement of YAP signaling^14^. HoxA1 is a common indicator of YAP activation in epithelial tissues, and was included to help determine YAP activation^36^. As YAP is normally expressed and sequestered in the cellular cytoplasm, mRNA levels in many cases will not change for YAP^37^. These genes as well as the NRG1 ligand were included in a qPCR panel of the same RNA extracts, and HoxA1 was found to be enriched at all time points. Because YAP signaling is most directly downstream of ErbB4 signaling, and enrichment was seen in both ErbB4 and HoxA1, this would suggest the importance of this pathway in the proliferative response observed.

**Figure 3:**
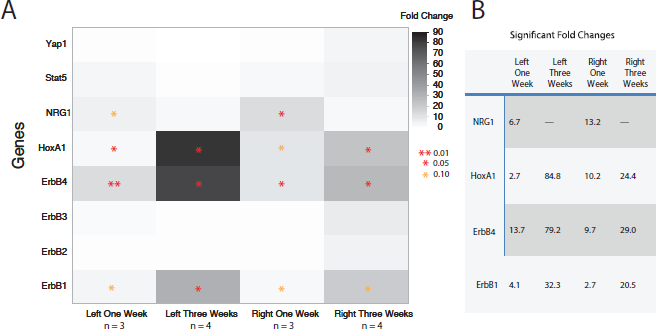
qPCR analysis of ErbB Receptor and downstream genes one and three weeks post amputation of the lung. Fig. 3. (A) Heat map of qPCR fold increases at both one wpa (n=3) and three wpa (n=4) for the left (injured) and right (contralateral) lungs. Red asterisks denote significant upregulation of mRNA products by two tailed T-test with unequal variance (p < 0.05), while orange asterisks denote trending towards significance (p < 0.10). B) Tables of significant fold changes (p < 0.05) for the qPCR results with fold changes indicated.

### Lineage Tracing Proliferation in the Lung after Injury

To determine the origin of cells during regeneration, animals were injected with 5-ethynyl-2′-deoxyuridine (EdU) two wpa, and either collected immediately (n=2), or at four wpa (n=4; Fig. 4A). Proliferating cells were analyzed at each time point and compared, both in the injured and contralateral lungs (Fig. 4B, C, D). Proliferation in the lungs was clustered, and each dividing cell only underwent approximately one division during the two-week chase period. The sources of the proliferating cells were lineage restricted, and epithelial cells that were dividing served to replenish epithelial layers in close proximity. Mesenchymal cells and ciliated cells also served to replenish local cells, staying within their own lineage. This pattern of proliferation suggests that there is not a specific stem cell niche in the axolotl lung that replenishes the entire tissue, but regeneration utilizes few divisions of many cells throughout the tissue to restore functional volume and mass. Mammalian lung regeneration is similarly lineage restricted ^38^. This stands as a dichotomy to limb regeneration in this species, where blastemal cells at the end of the amputated limb serve to replenish the cells of the entire new structure^21^.

**Figure 4:**
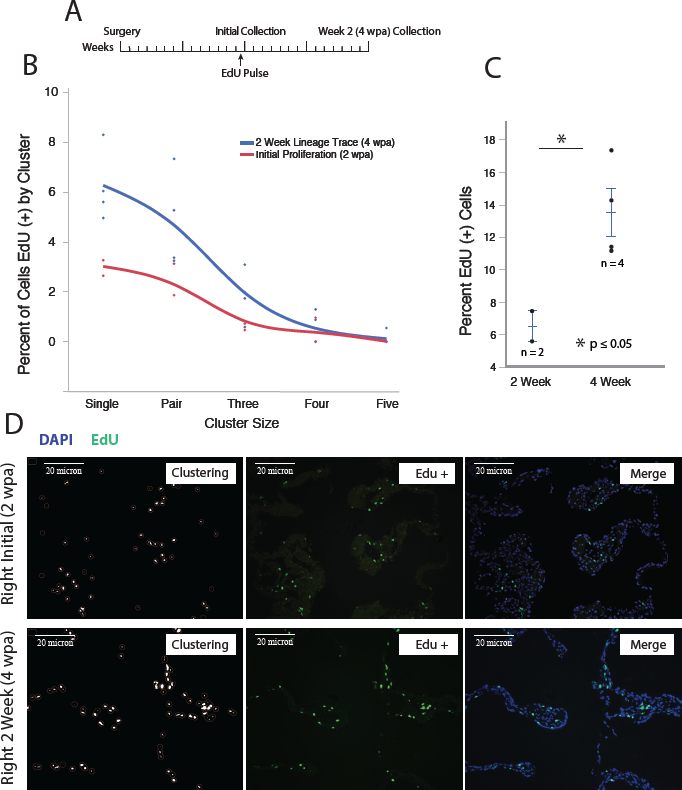
Lineage trace of lung tissue from two to four weeks post amputation. Fig. 4. (A) Representative experimental timeline of surgeries and collections with relative timing of EdU pulsing. Left lung was surgically injured in this experiment while right lung was left intact. Animals were collected at 2 wpa 3 hours after EdU pulsing (n=2) and animals were collected at 4 wpa, two weeks after EdU pulsing (n=4). Average measurements between the right and left lung were used. (B) Graph showing the distribution of the percent of dividing cells in each size cluster normalized to nuclear counts. Similar shape and distribution between the curves would indicate that the cells are only proliferating approximately once, and highly organized clusters are not developing. (C) Graph showing EdU (+) counts at initial (2 wpa) and 2 week chase (4 wpa). Two tailed T-Test with unequal variance shows significant (p = 0.03) upregulation of EdU (+) cells after the two week chase (4 wpa) (D) Example images of the clustering and counting used to measure the number of times cells would divide in the lineage labeled lungs. BioVoxxel toolkit for clustering in ImageJ was used to find the nearest neighbor within a 2-micron radius of each EdU+ cell, and cluster contiguous radii.

### Inhibition of ErbB2 Halts Proliferation

Mubritinib (TAK-165) is a highly specific (IC50 – 6 nM) inhibitor of the receptor ErbB2^39^. This receptor tyrosine kinase heterodimerizes with other members of the ErbB family and plays a key role in signal transduction of the ErbB receptors^9^. Because ErbB2 is not ligand binding, the heterodimer partner will provide specificity to the signaling molecule. Animals (n=4) were housed in water containing the ErbB2 inhibitor from days 12 through 21 dpa (Fig 5B,C). Animals showed no negative respiratory characteristics during treatment, such as increased gulping or pale appearance. In both injured and contralateral lungs, ErbB2 inhibition was sufficient to reduce cell proliferation approximately 4 fold, to similar levels seen in untreated animals. Barrier function of the lungs was unimpeded by the histological analysis of the tissue. Proliferation was reduced in both the injured and the contralateral lung, showing a global response to inhibition. ErbB2 plays an important role in NRG1β signaling, and its loss halts proliferation and regeneration in many tissue types^40,41^. It has been shown that inhibition of ErbB2 in this species prevents limb regeneration, and indicates a similar role for this family in controlling compensatory regenerative proliferation^42^.

**Figure 5:**
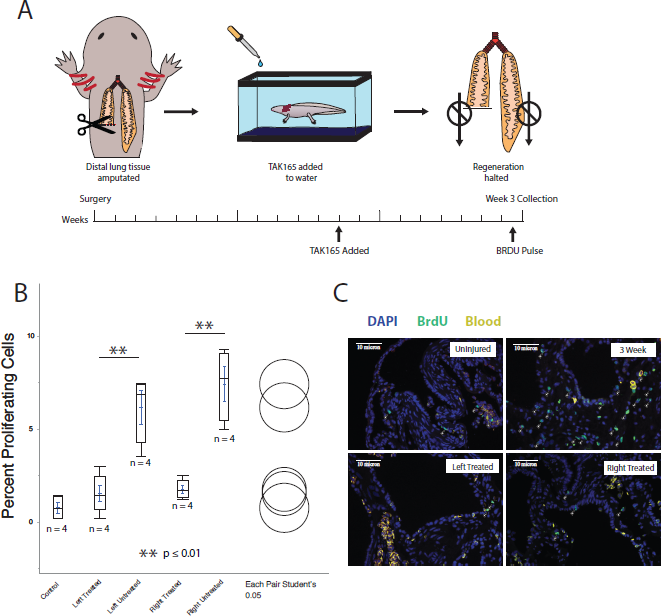
ErbB2 Inhibition halts lung regeneration by preventing cell proliferation. Fig. 5. (A) Representative experimental outline and timeline of surgeries and collections with relative timing of BrdU pulsing. Left lung was surgically injured in this experiment while right lung was left intact. Animals were treated with Mubritinib at 12 dpa and proliferation was measured at three weeks (n=4). This was compared to normal proliferation rates after injury at three weeks (n=4) and without injury (n=4). (B) Graph showing proliferation counts at control, normal left and right 3 wpa, and treated left and right 3 wpa,. Two tailed T-Test with unequal variance shows highly significant (Left p = 0.008, Right p = 0.007) downregulation of proliferation by the addition of TAK165 measured at three weeks. (C) Representative immunofluorescent images showing cell proliferation by anti-BrdU (Alexa488) antibody staining in control, normal 3 wpa, left treated 3 wpa, and right treated 3 wpa.

### Proliferation Induced by IP NRG1β Injection

NRG1 released by nerves has been shown to be necessary for limb regeneration in the axolotl and serves as a key ligand to ErbB4^42^. To understand the localization of NRG1 expression in the lung, we performed immunohistochemistry, and found strong staining on the apical surface of ciliated cells (Fig. 6A, B). To investigate other potential sources of NRG1 leading into the lung, we also discovered that the cranial nerve IX/X ganglion that innervates the lungs demonstrated had high expression levels of NRG1 (Fig. 6C). There are multiple potential sources of NRG1 in the lung, and further research must be performed to determine the contribution of various tissue types. The effect of NRG1β injection on whole axolotl lung tissue was investigated to ascertain whether NRG1β signaling would be mitogenic to lung tissue. Uninjured animals were injected with recombinant NRG1β IP (100 ng/g body weight). Proliferation was seen to increase significantly in the lung tissue, doubling the number of proliferating cells as compared to the sham-injected control (Fig 6D,F). Increased proliferation was observed throughout the affected tissue, and was not restricted to a specific lineage. This serves to highlight the importance NRG1β signaling in the lung tissue proliferative response.

**Figure 6:**
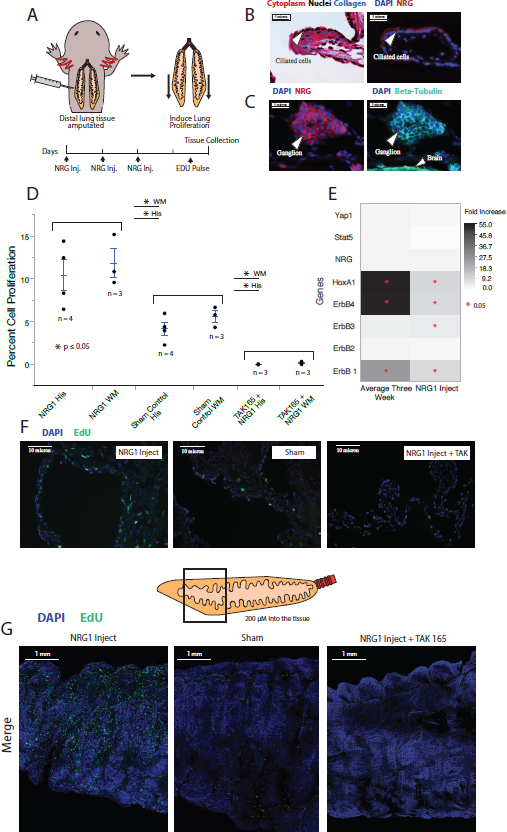
Proliferation induced in lung tissue by IP injection of Neuregulin-1. Fig. 6. (A) Representative experimental outline and timeline of injections and collections with relative timing of EdU pulsing. Lungs were left intact and NRG1 peptide was injected IP into animals (n=4). (B) Masson’s trichrome and NRG1 stained ciliated cells showing localization of NRG1 expression in the lung. (C) Co-staining of acetylated B-tubulin and NRG1 was present in the IX/X cranial ganglion, showing potential for axonal transport of NRG1. (D) Graph showing proliferation counts at sham control, after NRG injection and after NRG1 and TAK165 co-treatment, comparing whole mount and histology. Two tailed T-Test with unequal variance shows significant (Histology: p = 0.03 NRG1 injection, p = 0.01 NRG1 + TAK165; Whole Mount: NRG1 injection p = 0.05, NRG1 + TAK165 p = 0.01) upregulation of proliferation after NRG injection into to the animal versus water sham injection, downregulation of proliferation after TAK165 was added, and no significant difference in proliferation counts in histology versus whole mount. (E) Heat map of qPCRfold increases after injection of NRG (n=4) as compared to control (n=4). Red asterisks denote significant upregulation of mRNA products (p < 0.05 two tailed T-Test unequal variance). (F) Representative immunofluorescent images showing EdU+ cells in sham control NRG injection, and NRG injection while animals were treated with TAK165. (G) Whole mount lung representative images (n=3 per condition), merged channels Hoescht and EdU (green).

Additionally, qPCR was performed on lung tissue after NRG1β injection (Fig. 6E). It was found that similar pathways were upregulated to those seen in the injured tissues. Differences included a higher activation of EGFR than in injured lung tissue compared to uninjured, and some upregulation of ErbB3 in addition to ErbB4. It is likely that the greater upregulation of EGFR indicates an enhanced response of the tissue to cause differentiation of the new cells, especially in the absence of any injury or inflammatory signaling^43^. It has been previously studied that the interleukin IL-1B is important to NRG1β signaling, and its absence without injury could serve to impede signaling^17^.

NRG1β injection along with Mubritinib co-treatment led to inhibition of cell proliferation to levels observed in Mubritinib treatment alone (Fig. 6D,F) suggesting that ErbB2 is downstream of the proliferative response to the NRG1β injection. Mechanisms of proliferation in both regenerative and proliferative responses appear to be similar in nature. Animals tolerated to the co-dosage well and were visually inspected and deemed healthy. Lung tissue was examined by histology and appeared functionally normal aside from inhibition to lung proliferation. Altogether, this suggests that NRG1β signaling is mediated through ErbB family receptors, and these receptors appear to be vital for the induction of proliferation in pulmonary tissue.

### Whole mount visualization of treated lungs

To further visualize the proliferative response induced by NRG1β injection, a protocol was developed for whole mount lung tissue staining to observe cell proliferative responses throughout the tissue (Fig. 6G; supplemental material;). Animals were injected with NRG1β, co-treated with NRG1β and Mubritinib, or with sham control injections. No statistical difference was observed between whole mount data and histological sections, and allowed for a much greater number of cells to be analyzed in a shorter time.

## DISCUSSION

After traumatic lung injury, the lung must rapidly restore barrier function and gas exchange^44^. If significant tissue has been lost, it is vital that the functional volume and surface area of the lung be regained. In humans, there exists within the lung a niche of stem cells that are capable of restoring pulmonary tissue, but recovery is very slow^6^. Studying model organisms that can rapidly regenerate can serve to help us uncover mechanisms and signaling molecules through which human lung tissue may be induced to proliferate and restore efficient lung function.

This study provides evidence that the axolotl salamander is capable of significant regeneration of the lung after amputation. After the distal third of the lung was removed, the tissue rapidly closed off the injury site, restoring the barrier function of the epithelium and resisting scar formation. Lung cells began dividing and pulmonary tissue grew larger to compensate for what had been lost. While it did not appear that were large differences in the proliferation rate throughout the organ system, after eight weeks the tissue in the injured lung had caught up to the contralateral lung. While the proliferation and mass increased in the injured lung, it did not recover the length seen in the contralateral lung. Interestingly, lung growth was not directional proximal to distal and instead occurred in all directions outward to expand the lung volume. During salamander limb regeneration, proliferation is witnessed within close proximity to the injury site, and contributes to the formation of a blastema. It appears that lung regeneration is mechanistically distinct from the regenerative response that occurs in the limb.

It was found that there was significant upregulation of receptors ErbB4 and EGFR at both one week and three weeks after distal amputation in the lung, as well as an upregulation of the signaling ligand NRG1β. EGFR has been indicated in promoting lung cell maturation and differentiation^34^, and a ligand of ErbB4, NRG1β, has been shown to play an important role in lung development^45,46^. When used to activate ErbB4, NRG1β can control both lung epithelial cell proliferation and surfactant synthesis in vitro in mammalian pulmonary cells^30^. In our experiments, inhibition of ErbB2 reduced cell proliferation in both the injured and the contralateral lung, showing a global response to inhibition. ErbB2 plays an important role in NRG1β signaling, and its loss halts proliferation and regeneration in many tissue types^28,29,41^. This would indicate that NRG1β signaling to ErbB4:ErbB2 heterodimers is most likely controlling pulmonary cell proliferation during regeneration. In human patients after ALI, physicians have noted elevated levels NRG1 in bronchoalveolar lavage^47^. This would lead to the conclusion that this pathway is present in human lung tissue, and holds potential for therapeutic intervention, warranting further study to fully elucidate the potential of NRG1 as a therapeutic pneumocyte growth factor.

It is also known that once ErbB4 is activated, it can undergo proteolytic cleavage and release its intracellular domain^16^. This domain, can serve to shuttle Stat5 to the nucleus^35^, and once in the nucleus, it can enhance Yap signaling, upregulating many developmental and proliferative genes^48^. A gene that has been used as an indicator of Yap activation in cancers is HoxA1^36^. HoxA1 was seen to be upregulated at one week, with enrichment continuing into 3 weeks post amputation. While Yap was not upregulated, this is not unusual as it is often sequestered in the cytoplasm ready for release, and does not need to increase in expression to confer proliferative affects^37^. Yap activation has a potential as a potent downstream effector of regeneration^49^.

We have demonstrated that exogenous IP injection of NRG1β peptide is sufficient to recapitulate the response seen after injury, and upregulate proliferation in the lung tissue. There has been much conjecture over the years as to the identity of the pneumocyte growth factor, and we have provided evidence that NRG1β may be this molecule^12^. Transcript profiles between injured tissue and post NRG1β injection tissues were similar, with the key difference being EGFR being more highly enriched. This is likely due to the absence of inflammatory response, and the tissue exhibiting an increased propensity towards differentiation in the intact tissue. As NRG1β does not signal the EGF Receptor, this is the most likely explanation. As was seen in lung regeneration, the proliferative response to NRG1β was blocked by ErbB2 inhibition. To visualize global proliferation, we developed a rapid protocol to visualize cell proliferation in whole mount axolotl lungs utilizing EdU Click-it technology. This is the first time we have seen this technique utilized, and should be a useful technique for studying changes in cell proliferation across entire organs in other systems. We have provided the first glimpse at how this model organism regenerates its lung tissue, and provided a basis for further research into this species’ lung regeneration.

## CONCLUSIONS

In this study, we have shown that Axolotl lung tissue regenerates using a compensatory mechanism, contrary to the epimorphic limb regeneration observed in this species. Epidermal growth factor signaling is crucial for regeneration to take place, and appears to be specifically through ErbB2:ErbB4 heterodimer receptors. Neuregulin-1 can induce proliferation in the lung, and is a likely candidate to exert molecular control over lung regeneration. ErbB4 could hold therapeutic value for future research, and further studies in this species could provide novel insight into mechanisms through which mammalian lung regeneration may be enhanced after injury.

## METHODS

### Animal Use and Study Design

IACUC of Northeastern University approved this study under protocol number 15-1138R. All experimental procedures and animal care were conducted in accordance to vertebrate care guidelines. Animals were on average 13 cm in length and 7 cm in snout to vent length, six months old, and were raised in Northeastern university lab facilities according to Farkas and Monaghan, 2015. Animals were kept in individual tanks with regular water changes and fed three times a week. Sample size was selected after seeing a large effect size in preliminary data, justifying small n values. No exclusion criteria were determined; all animals were included. Animals were non-randomly assigned to groups to ensure all animals were at the same stage of development. No blinding was performed.

### Surgical Procedures

Axolotls were sedated by immersion in 0.01% benzocaine solution. An incision was made above the spleen according to Fig. 1A. Forceps were then inserted into the abdominal cavity through the small hole, passing beneath dorsal muscles running parallel to the spine. The distal lung tip was pulled through the incision, and a third of the lung was amputated using dissecting scissors. Forceps were then used to push the remaining lung tissue away from the incision, so as to prevent tissue adhesion at the wound site. The wound was then closed with 3M Vet Bond tissue adhesive. The axolotl was placed back into animal housing, with daily observation to check recovery progress.

### Tissue Processing and Histology

The flank of euthanized animals was opened using dissecting scissors and right and left lungs removed. Insulin syringes were used to inflate lungs with 10% neutral buffered formalin (NBF) while forceps were used to seal the bronchial openings. Lungs were then submerged in NBF and fixed overnight at 4°C. After fixative treatment, lungs were washed in phosphate buffered saline 3x and immersion in 70% ethanol. Tissues were processed for paraffin embedding and sectioned to 8-micron sections. Slides were heated at 55°C for one hour to adhere wax sections to the slides prior to deparaffinization and staining. Tissues underwent Masson’s Trichrome Straining (Thermo Scientific Chromaview) for visualization of different cell types. For visualization and staining of pneumocytes, sections were rehydrated and then held in BM Purple (Roche) overnight at 4°C and stained with eosin to differentiate pneumocytes from surrounding tissue.

### Cell Proliferation and Immunohistochemistry

Animals were anesthetized in 0.01% benzocaine and IP injected with BrdU at 1 mg/g or EdU 25 ng/g in saline by body mass 12 hours or 3 hours prior to euthanization and collection, respectively. Histology was performed as previously described in Farkas et al., 2016. Primary antibodies (Krt7 1:500, BrdU 1:500, NRG 1:1000) were diluted in goat serum + PBS and placed on blocked sections incubated overnight at 4°C. Secondary antibodies were diluted in PBS and incubated on sections for 30 minutes (1:500). Sections from EdU-pulsed animals were deparaffinized and placed in EdU reaction mixture as listed in supplement materials for 30 minutes at room temp. DAPI nuclear stain was then added and slides were mounted and imaged.

### Whole Mount Preparation

Lungs were extracted after EdU injection and inflated with 4% PFA prior to submersion overnight at 4°C. Tissues were then dehydrated and rehydrated through a methanol/PBS series and permeabilized with trypsin prior to staining with FAM-azide conjugation to EdU. Tissues were then submerged in 70% glycerol with Hoescht overnight at 4° C. Tissues were placed in a new wash of 70% glycerol and imaged using laser-scanning confocal microscopy. Stacks were taken through approximately 200 micron of the tissue and Z-stack projections were generated using Image J. Whole mount protocol is further described in the supplementary material.

### Drug Treatment

Mubritinib (TAK 165) (TSZ Scientific) stock solution (10mM in DMSO) was diluted in salamander housing solution to 1 μM. Animals were treated at 12 dpa and collected at three wpa. Animals were pulsed with BrdU as previously described to measure proliferation rates in the treated animals. Animals were euthanized and lungs collected 12 hours post BrdU injection for immunohistochemistry.

### NRG1 Injection

Animals were anesthetized in 0.01% benzocaine and NRG-1 was injected at a concentration of 100 ng of recombinant human NRG1β-1 peptide per gram animal weight per day, for three days (Peprotech, 100-03). At three days post-treatment animals were injected IP with EdU, euthanized 3 hours later, and lungs collected for sectioning and mounting.

### qPCR Analysis

Lungs were collected from animals at one and three wpa, and lungs were flash frozen using liquid nitrogen and stored at -80 °C. Total RNA was extracted using TRIzol Reagent (Life Technologies) followed by Qiagen RNeasy kits according to manufacturer’s protocol. Samples were transcribed to cDNA using Verso cDNA Synthesis Kit (Thermo Scientific). qPCR was performed using SYBR Green Supermix (Applied Biosystems), cDNA according to 25ng of total RNA, and 0.5μM of each primer. qPCR was performed with paired technical replicates and with biological replicates of three or four as listed. Expression levels for genes were normalized using β-actin as a control gene. Primers were made using Primer 3 software and axolotl transcriptomics data courtesy of axolotlomics.org. qPCR was performed in a Step One qPCR system (Bio-rad). Relative messenger RNA expressions were calculated using the 2^-ΔΔCT^ method. The following primers were used for amplification:

F_YAP 1 _Isofo rm_3: 5′-TGTTCCCAGAACACCAGATG-3′;

R_YAP1_Isoform_3: 5′-GTAATCTGGGAAGCGGGTTT-3′;

F_Hoxa1: 5′-GCTGGAGAGTACGGATACGC-3′;

R_Hoxa1: 5′-TGGAACTCCTTCTCCAGCTC-3′;

F_Stat5: 5′-CCGGAGCAAGTTACATGGAT-3′;

R_Stat5: 5′-TCAGGGTCCAGAATGGAGTC-3′;

F_Erbb4: 5′-CGCAGGCCAGTCTATGTAAT-3′;

R_Erbb4: 5′-TTAGTGGCTGAGAGGTTGGT-3′;

F_Erbb2: 5′-GGAACTTCTCCCCAGTATCC-3′;

R_Erbb2: 5′-CATGGAGGGTCTTTGATACC-3′;

F_Egfr: 5′-GCCAAGTGAAACCAAAGTCC-3′;

R_Egfr: 5′-CTTGGCGTGTTCTGGTATTC-3′;

F_Erbb3: 5′-GCTACTGAACTCGGTGAGTG-3′;

R_Erbb3: 5′-GTCGGATCAGAGCTGTACCT-3′;

F_Nrg1: 5′-CGAGTGCTTTGTCCTCAAG-3′;

R_Nrg1: 5′-CAGCGATCACCAGTAAACTC-3′.

F_B Actin: 5′-AGAGGGGCTACAGCTTCACA-3′

R_B Actin: 5′- GGAACCTCTCGTTGCCAATA-3′

### Statistical Analysis

JMP12 (SAS Institute Inc.) was used for data analysis. Data analysis was performed by calculating each pair two tailed unequal variance student’s T-Test to test for significance; p ≤ 0.10 was considered trending significant, p ≤ 0.05 was considered significant and p ≤ 0.01 was considered highly significant. All error bars represent SEM and center lines represent mean values.

## DECLARATIONS

### Ethics approval and consent to participate

All experiments performed in accordance with IACUC protocols and in alignment with departmental regulations. IACUC of Northeastern University approved this study under protocol number 15-1138R.

### Consent for publication

Not applicable

### Availability of data and materials

All data is available from the author upon request.

## Authors' contributions

TJ designed the experiments, conducted the experiments, analyzed the data and wrote the manuscript, providing intellectual leadership from conception to finalization. PG and NGS conducted experiments and analyzed data. YK, HW, AA conducted experiments. JRM supervised work, designed experiments, and contributed to the data analysis and publication formulation, as well as provided mentorship to TBJ. All authors contributed to editing the manuscript.

## Acknowledgements

Thanks to Johanna Farkas for her work in adapting the whole mount procedure, and thanks to Alex Lovely for his work performing the confocal imaging of the whole mounted lung tissue.

Thanks to the Andrew I Schafer Co-op Scholarship for salary funding.

Thanks to the Northeastern University Biochemistry Program for financial support.

## Authors' information

Article written and prepared as senior thesis of TBJ. JRM provided mentorship and support. Correspondence and requests for materials may be addressed to JRM.

